# Probing drug-target engagement of soluble Gα_i1_ protein using the SolThermoBRET thermal shift assay

**DOI:** 10.1101/2024.02.15.580002

**Authors:** Oscar Brod, Darya Plevako, Themiya Perera, Pattarin Hompluem, Eline J. Koers, Manuel Hilbert, David A. Sykes, Dmitry B. Veprintsev

## Abstract

**Introduction:** Sensitive protein stability assays used during hit confirmation conventionally require high quantities of purified protein. Here, we describe a novel high-throughput 384-well BRET-based thermostability assay allowing for the ultrasensitive determination of Gα_i1_ protein stability. Using this method we make use of the environmentally sensitive dye, SYPRO Red, which interacts with hydrophobic regions in the protein that become exposed upon denaturation. Assays are functional in crude cell lysates, without any requirement for protein purification.

**Methods:** The SolThermoBRET method measures resonance energy transfer between a thermally stabilised nano-luciferase (tsNLuc) genetically engineered to the N-terminus of Gα_i1_ protein, and SYPRO Red, a fluorescent dye that binds to lipophilic residues exposed upon protein unfolding in response to thermal denaturation. Samples in 384-well PCR plate were subjected to a temperature gradient (30 – 60 °C) on a PCR block and incubated for 5 minutes. Isothermal SolThermoBRET at a fixed temperature 45°C allowed the measurement of IC_50_ curves to various ligands. Furimazine (10 μM) was added to the samples and BRET read on the BMG Labtech PHERAstar FSX at room temperature. Melt curves were fit to a Boltzmann sigmoidal equation to obtain Tm values.

**Results:** Ligand-induced stabilisation of Gα_i1_ protein was demonstrated in non-purified samples. A range of Tm values were measured with increasing stabilisation observed for nucleotides/sides in the following order GDP<GTP<GppNHp<GTPγS. Correlation was observed between SolThermoBRET derived Tm’s determined in the presence of high ligand concentrations and IC_50_ determinations using Isothermal SolThermoBRET.

**Conclusions:** SolThermoBRET represents a sensitive nanoscale method for the detection of changes in soluble protein thermostability. This method is attractive for hit conformation and the identification of novel ligands targeting soluble proteins.

## Introduction

The process of developing novel therapeutics requires an assay that directly detects target engagement to validate the results of any functional high-throughput screen [1]. Thermal shift assays (TSA) have emerged as a pivotal tool in this endeavour, as the stabilisation of proteins by ligands follows the fundamental laws of physical chemistry and is a universally observed phenomena. The key advantage of methods such as TSA is that they do not depend on a target specific tracer to assess the potential of ligands to bind a particular target. TSA methods have the potential to assess target engagement in the absence of known ligands, and together with universal applicability this makes them especially powerful tools to assess previously unexplored drug targets.

A plethora of biophysical methods have been established in the last 50 years to measure protein stability, including differential scanning calorimetry (DSC), circular dichroism spectroscopy (CD) [2], intrinsic protein fluorescence and light scattering. In case of soluble proteins (ca. 75% of the human proteome), differential scanning fluorimetry (DSF) methods utilising environmentally sensitive fluorescence dyes such as anilinonaphthalene sulfonate (ANS) and Sypro™ Orange and Sypro™ Red (called SYPRO Red thereafter), have emerged as a particularly rapid and accessible method to unravel their stability landscape and stabilisation by ligands [3-11].

In the early phases of drug discovery, establishing over-expression systems and purification of the target protein itself can pose significant challenges, slowing down the discovery process. Indeed, one major drawback of traditional TSA is that they require a significant concentration of purified, detergent free protein (∼ μM) to establish a good signal-to-noise. This limits their applicability in cases when purifying large amounts of the target protein in the functional form is difficult, or if there are unknown co-factors that are important for stability or function.

Several methods have been developed to study protein stability and their stabilisation by ligands in the cellular or cell lysate setting, addressing some of the limitations of the biophysical methods mentioned above. These include limited proteolysis [12] and mass-spectrometry [13, 14] which broadens up the scope of the limited proteolysis method to the whole proteome. CETSA (cellular thermal shift assay) is another popular technology that monitors the binding of ligands via changes in protein aggregation and does not require modification of the protein target, with the added advantage that it can be applied in living cells [15]. However, CETSA often depends on the availability of specific antibodies which limits the number of samples that can be processed, as well as the targets which can be interrogated [16]. Recently, a more universal detection approach has been introduced using a split NanoLuc to alleviate the need for specific antibodies [17]. Both DSF and CETSA methods have proved advantageous for high throughput screening in an industrial setting [18].

However, it would be advantageous to combine the accuracy and convenience of the DSF screening method, employing purified proteins with the ability of limited proteolysis and CETSA to monitor the stability of unpurified protein in cell lysates. To address the critical need for an ultrasensitive thermostability assay applicable to non-purified soluble proteins, we adapted the ThermoBRET approach that we previously developed to measure the stability of membrane proteins, [19] and applied it to soluble proteins. SolThermoBRET monitors changes in BRET between a thermally stabilised nano-luciferase (tsNLuc) [20] tagged protein and SYPRO Red, a fluorescent dye that binds hydrophobic surfaces exposed as a soluble protein undergoes thermal denaturation. In the case of SolThermoBRET, the fluorescent properties of the dye SYPRO Red are better suited for excitation by the tsNLuc protein due to its lower reported background signal.

As a proof of concept, we applied this method to the Gα_i1 i_ protein which plays a crucial role in cell signalling, transmitting chemical information via GPCRs on the outside of the cell to the inside, in a process known as signal transduction. The Gα_i1_ protein is primarily involved in inhibiting signalling transduction, turning off cAMP production inside the cell by inhibiting adenylyl cyclase activity. In the native environment of the cell these enzymatic proteins, with inherent GTPase activity, bind to their substrates the guanine nucleotides, guanine diphosphate (GDP) and guanine triphosphate (GTP) [21]. Due to its abundance in all cells, inhibitors of this protein are unlikely to make good therapeutics, however experimental tool compounds may have some use in disease models where excessive Gα_i1_ signalling is thought to play a role.

SolThermoBRET presents several advantages compared to traditional TSA. It demands a lower protein concentration, reduces sample preparation times, and eliminates the need for protein purification. This lowering of barriers allows for the study of proteins with limited availability and potentially facilitates its application in high-throughput scenarios.

## Material and methods

### Materials

Dulbecco′s Phosphate Buffered Saline (without calcium chloride and magnesium chloride, Cat #D8537-500mL) Glycerol (Cat #49770-2.5L), BSA (Cat #A7030-100G) and DL-Dithiothreitol (DTT) (Cat #D0632), Adenosine 5′-diphosphate (ADP #O1905), Guanosine 5′-diphosphate sodium salt (GDP #G7127), Guanosine 5′-triphosphate sodium salt hydrate (GTP #G8877), Guanosine 5′-[β,γ-imido]triphosphate trisodium salt hydrate (GppNHp #G0635) and Guanosine 5′-[γ-thio]triphosphate tetralithium salt (GTPγS #G8634) were obtained from Sigma-Aldrich. Furimazine (or NanoGlo substrate) was obtained from Promega (Cat #N2012). FrameStar® 384 Well Skirted PCR plates (#4ti-0385) were obtained from Scientific Laboratory Supplies (SLS) and 96 well Semi-skirted LP PCR plates, white (#LW2230W-10) were obtained from Alpha Laboratories. SYPRO Red Protein Gel Stain (5,000X Concentrate in DMSO) was obtained from Invitrogen Molecular Probes.

## Methods

### Gα_i1_ protein production

The tsNLuc [20] was inserted into the human 10His-Gα_i1_ construct in pJ411 bacterial expression vector described earlier[9] between the his-tag and N terminus of the Gα_i1_. Sequence was confirmed by Sanger sequencing. NiCo21(DE3) chemically competent E. coli were transformed with pJ411 bacterial expression plasmids and plated onto LB/agar plates containing 2% w/v glucose and 50 μg/mL kanamycin. After incubation at 37 ºC for 16-24 hours, a single colony was picked to inoculate 20 mL of terrific broth containing 0.2% w/v glucose and 50 μg/mL kanamycin. After 16-24 hours in a shaking incubator set at 37 °C, 15 mL of overnight culture was added to 3 L of terrific broth containing 0.2% w/v glucose and 50 μg/mL kanamycin, grown in a shaking incubator at 37 °C until OD_600_ of 0.7, when 1 mM of isopropyl-β-D-thiogalactopyranoside (IPTG; VWR Chemicals) was added to induce protein expression. Cells were then grown overnight (16-20 hours) at 23 °C in a shaking incubator before being harvested by centrifugation and frozen at -80 °C. Cell pellets were then thawed on ice, and resuspended in 30 mL (per 800 ml of bacterial culture) lysis buffer (25 mM phosphate buffer, 150 mM NaCl, 10% glycerol, 5 mM DTT, pH 7.2 supplemented with 0.25 mg/mL chicken lysozyme, 1 μg/mL bovine DNAse I, and cOmplete™ EDTA-free Protease Inhibitor cocktail tablets (Roche)). Cells were then lysed on ice by sonication (). Cell lysates were then clarified by centrifugation at 25,000 rcf for 30 minutes and then by passing through a 0.45 μm syringe filter. The filtered lysate was aliquoted, frozen in liquid N_2_ and stored at -80 °C until use.

### Gα_i1_ protein titration

Assays were performed in Dulbecco′s Phosphate Buffered Saline (without calcium chloride and magnesium chloride), liquid, sterile-filtered, suitable for cell culture and containing 10% Glycerol 0.01% BSA plus 5mM DTT. Unpurified Gα_i1_ protein was serially diluted in assay buffer using a 2-fold dilution steps from an initial dilution of 1000 to 64,000. Diluted samples (10 μL) of unpurified Gα_i_ protein in assay buffer were then dispensed in a white 384-well PCR plate. Furimazine (10 μM) was then added to the samples and the plate read on the BMG Labtech PHERAstar FSX at room temperature.

### SolThermoBRET on unpurified Gα_i1_ protein

#### Thermal shift experiments

Assays were performed in Dulbecco′s Phosphate Buffered Saline (without calcium chloride and magnesium chloride), liquid, sterile-filtered, suitable for cell culture and containing 10% Glycerol 0.01% BSA plus 5mM DTT. Nucleotide/side were initially dissolved in water and intermediate dilutions prepared in assay buffer. Samples (10 μM L) of unpurified Gα_i1_ protein (diluted 32,000-fold) in assay buffer containing either vehicle or nucleotide/side (100 μM) were dispensed in a white 384-well PCR plate were subjected to a temperature gradient (30 – 60 °C) on a PCR block and incubated for 5 minutes. Furimazine (10 μM) was then added to the samples and the plate read on the BMG Labtech PHERAstar FSX at room temperature.

#### Isothermal SolThermoBRET

Isothermal SolThermoBRET at a fixed temperature 45°C allowed the measurements of IC_50_ curves for various ligands. Assays were performed in Dulbecco′s Phosphate Buffered Saline (without calcium chloride and magnesium chloride), liquid, sterile-filtered, suitable for cell culture and containing 10% Glycerol 0.01% BSA plus 5mM DTT. Samples (10 μL) of unpurified Gα_i1_ protein (diluted 32,000-fold) in assay buffer containing either vehicle or nucleotide/side (100 μM) were dispensed in a white 384-well PCR plate containing 1uL of nucleotide/side or vehicle and were subjected incubated for 5 minutes on a PCR block at a fixed temperature 45°C. Furimazine (10 μM) was added to the samples and read on the BMG Labtech PHERAstar FSX at room temperature.

### Data analysis

For luminescence thermostability measurements, unfiltered luminescence was normalised to the top point of the dataset. SolThermoBRET melt curves were fit to a Boltzmann sigmoidal equation (see equation, Eq. 1). This curve-fitting procedure was performed using GraphPad Prism 10 and employs a sigmoidal function to fit the protein thermal denaturation curve, such that T_m_ can be determined by the inflection point.

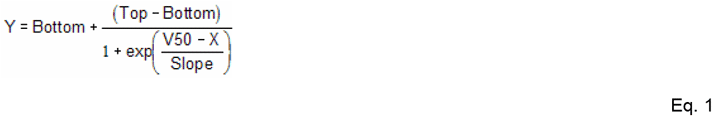

This equation conventional describes voltage dependent activation of ion channels. Here it describes the degree of protein unfolding (Y) as a function of the temperature (X). Unfolding varies from BOTTOM to TOP. V50 in this case is the temperature (or T_m_) at which protein unfolding is halfway between BOTTOM and TOP.

For SolThermoBRET measurements, NanoBRET ratio was defined as the 610LP emission divided by the 450BP80 emission. The data was normalised to the upper (100% unfolded) and lower (0%) datapoints and fitted using a Boltzmann sigmoidal equation. 100% defined as the degree of unfolding observed in the presence of GTP.

For Isothermal SolThermoBRET experiments data was fitted to a four parameter logistic equation (see equation, Eq. 2) to obtain IC_50_ estimates.

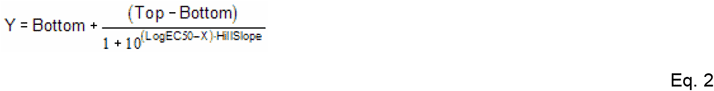

Where, Bottom is the Y value at the bottom plateau. Top is the Y value at the top plateau. IC50 (inhibitory concentration, 50%) is the X value when the response is halfway between Bottom and Top. And the HillSlope describes the steepness of the curve.

## Results

ThermoBRET applied to soluble intracellular proteins, employs a combination of advanced techniques, namely Bioluminescence Resonance Energy Transfer (BRET) and environmentally sensitive dyes such as SYPRO Red, to assess the thermostability of a soluble protein labelled with tsNLuc, in this case the Gα_i1_ protein.

### Optimization of the Gα_i1_ protein concentration

To determine a suitable concentration of Gα_i1_ protein for testing in the SolThermoBRET assay it was necessary to check the luminescent levels at various dilutions of the unpurified protein. The goal was to obtain sufficient bioluminescence signal that is detectable after ca 15-20 min. Too much of the tsNLuc -tagged protein would deplete the substrate (furimazine) too quickly. Optimisation of Gα_i1_ protein concentration is shown in Figure 2, along with a cartoon illustrating the relatively low protein requirements of SolThermoBRET versus other biophysical techniques.

**Figure 1.**
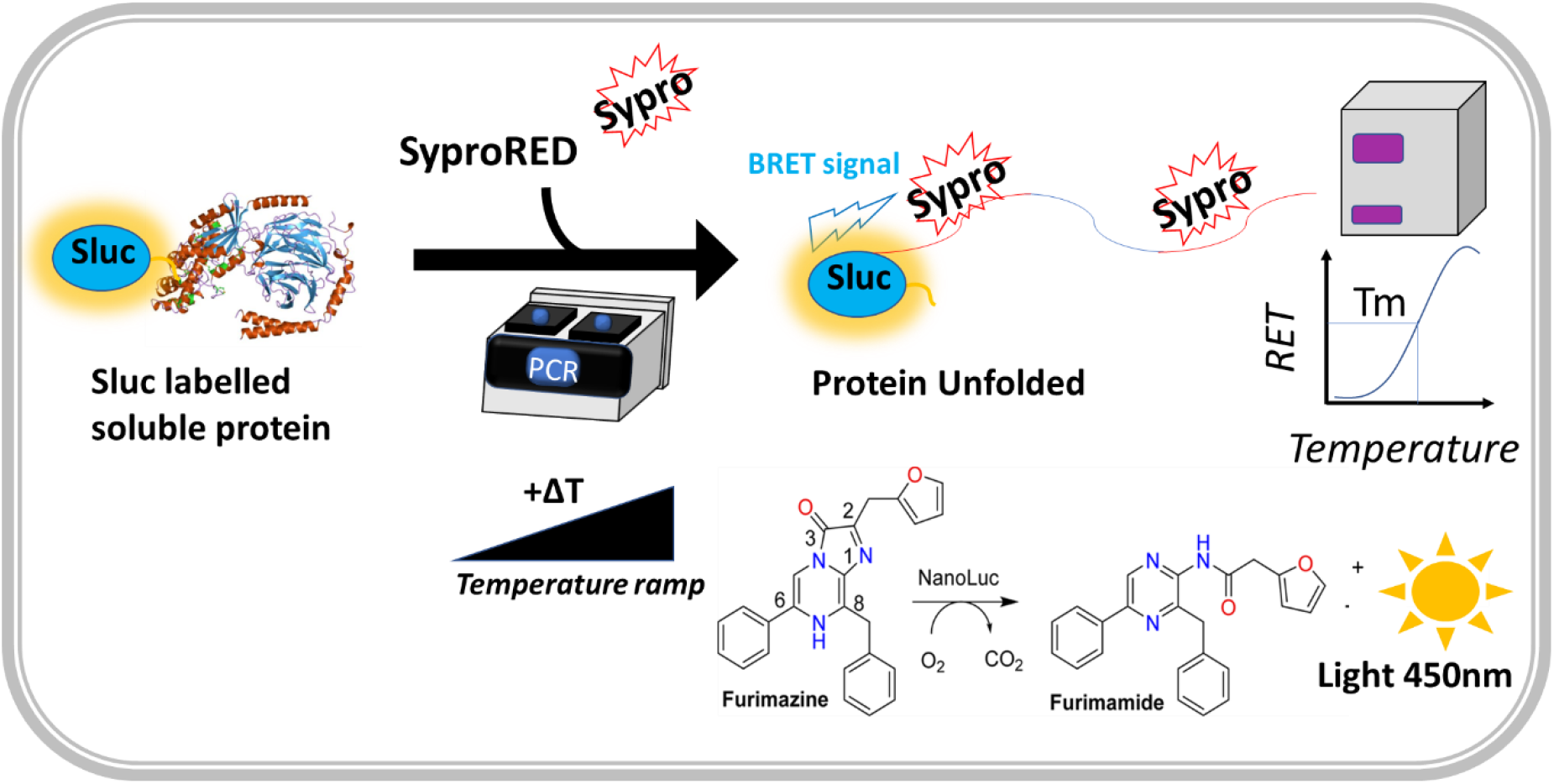
SolThermoBRET thermostability assay principles. The environmentally sensitive dye SYPRO Red interacts with hydrophobic regions in the protein exposed upon denaturation. This interaction is detected via BRET between the tsNLuc protein and the dye in proximity.

**Figure 2.**
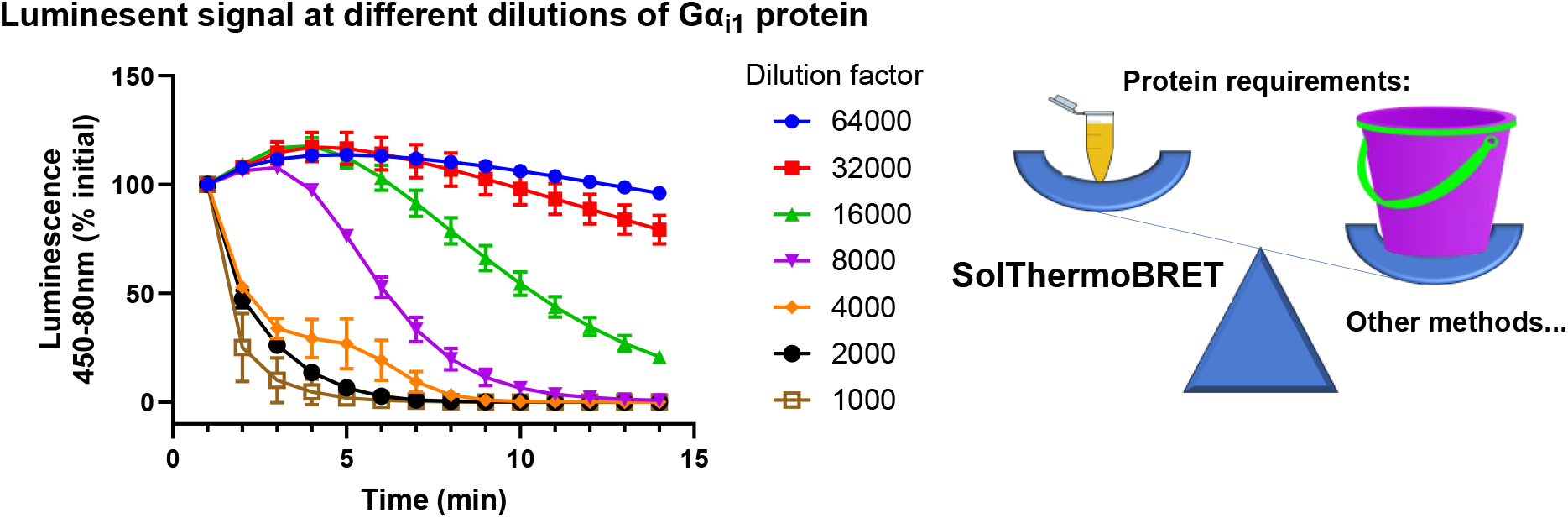
Optimisation of Gα_i_ protein concentration and relative protein requirements of SolThermoBRET versus other biophysical techniques. Data are shown as mean ±standard error (n≥3).

### Ligand-induced stabilization of Gα_i1_ protein

Ligand-induced stabilisation of Gα_i1_ protein was demonstrated in non-purified samples by following the unfolding of the protein in the presence of different nucleotides/sides and the environmentally sensitive dye SYPRO Red.

The effect of high concentrations of various nucleotides/sides on Gα_i1_ protein thermostability relative to the apo protein are shown in Figure 3. Ligand induced thermostability shifts were clearly visible in the following order from low to high Tm values as follows; GDP<GTP<GppNHp<GTPγS, see Table 1. ADP gave no shift suggesting that either it does not bind to Gα_i1_ protein, or it has an affinity lower than the 100 μM employed in these assays.

**Table 1.** T_m_ measures in the presence of various nucleotides/sides relative to apo protein. Data shown as mean ±standard error (n≥3).

| Nucleotide/side | T<sub>m</sub> °C (n) |
| --- | --- |
| Control | 42.96 ± 0.63 (4) |
| GDP | 46.64 ± 1.31 (3) |
| GTP | 48.33 ± 0.44 (4) |
| GppNHp | 50.42 ± 0.32 (4) |
| GTP<sub>γ</sub>S | 52.94 ± 1.46 (4) |
| ADP | 43.14 ± 0.54 (4) |

**Figure 3.**
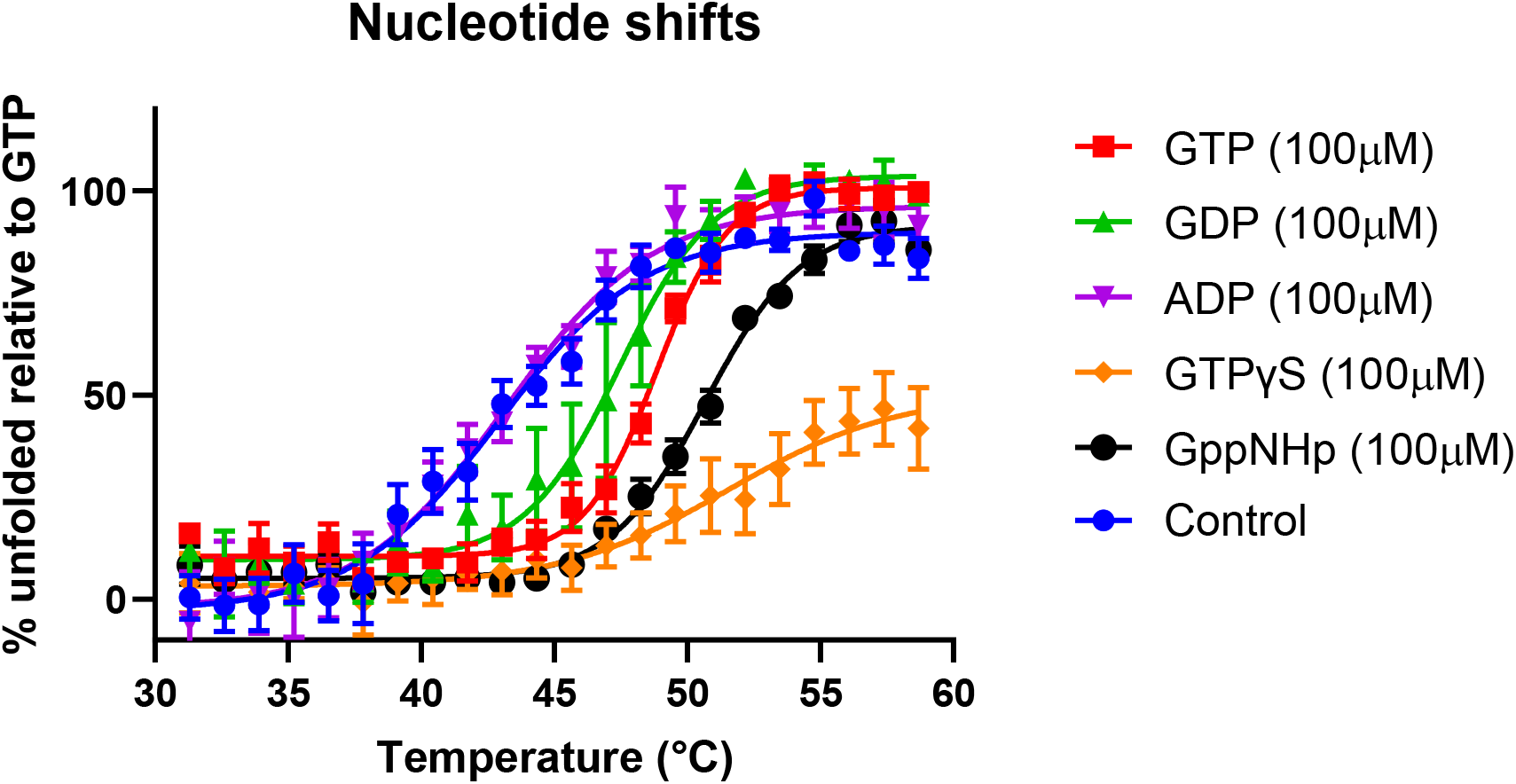
SolThermoBRET ligand thermostability shifts. Effect of high concentrations of various nucleotides/sides on Gα_i1_ protein thermostability relative to the apo protein. Data are shown as mean ±standard error (n≥3).

### Isothermal SolThermoBRET experiments

Isothermal SolThermoBRET experiments were then performed to measure the relative affinity of the different nucleotides/sides tested in the SolThermoBRET thermal shift assay. Inhibition of protein unfolding at a constant temperature in the presence of various concentrations of the four nucleotides/sides which shifted the melting curve is shown in Figure 4A-D. Concentration dependent, ligand induced inhibition of Gα_i1_ protein unfolding was observed. Moving from low to high affinity as follows, GDP<GTP<GppNHp<GTPγS. IC_50_ values, which represent the concentration of ligands at which half-maximal inhibition of protein unfolding occurs, were determined from the isothermal SolThermoBRET experiments and are reported in Table 2.

**Table 2.** IC_50_ and delta T_m_ measures of various nucleotides/sides. Data shown as mean.

| Nucleotide/side | pIC<sub>50</sub> (n) | Delta T<sub>m</sub> (n) |
| --- | --- | --- |
| GDP | 5.45 ± 0.31 (5) | 3.87 ± 1.93 (3) |
| GTP | 5.76 ± 0.20 (6) | 5.37 ± 0.98 (4) |
| GppNHp | 6.12 ± 0.27 (6) | 7.46 ± 0.53 (4) |
| GTP<sub>γ</sub>S | 6.77 ± 0.29 (6) | 9.98 ± 1.40 (4) |
| ADP | ND | 0.18 ± 0.71 (4) |

**Figure 4.**
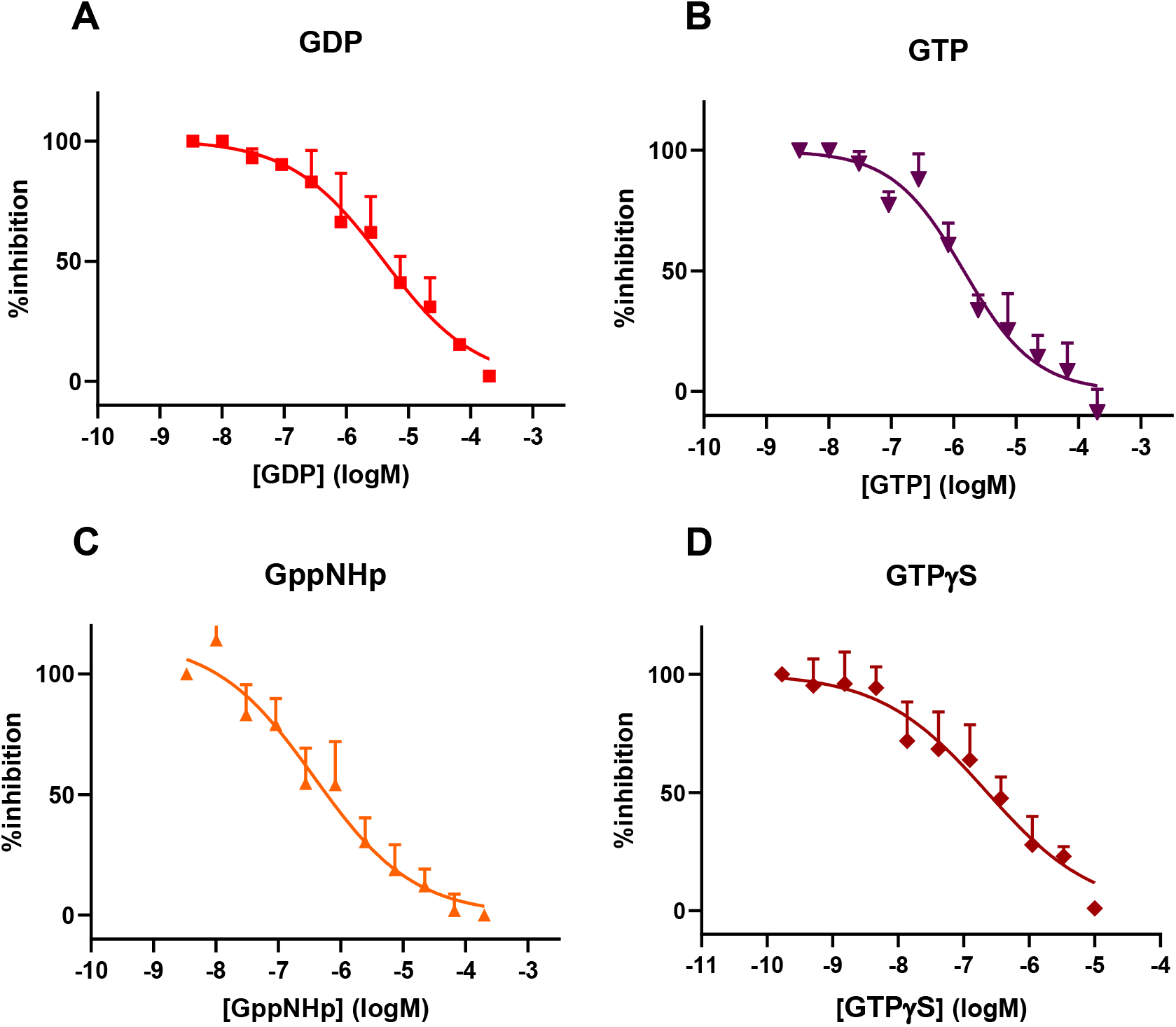
Isothermal SolThermoBRET. Inhibition of Gα_i1_ protein unfolding at a constant temperature in the presence of various concentrations of **A)** GDP **B)** GTP **C)** GppNHp and **D)** GTPγS. Data are shown as mean ±standard error (n≥3).

### Correlation between ΔTm and IC_50_ parameters

To better understand the relationship between the two different assay formats, we plotted the SolThermoBRET derived changes in melting temperature (ΔT_m_) determined in the presence of high ligand concentrations and the IC_50_ determinations obtained using the isothermal SolThermoBRET method. An almost perfect correlation was observed between the two measures suggesting that the degree of shift observed in the SolThermoBRET assay is related to the relative affinities of the ligands for the target and the concentrations tested, see Figure 5 and Table 2.

**Figure 5.**
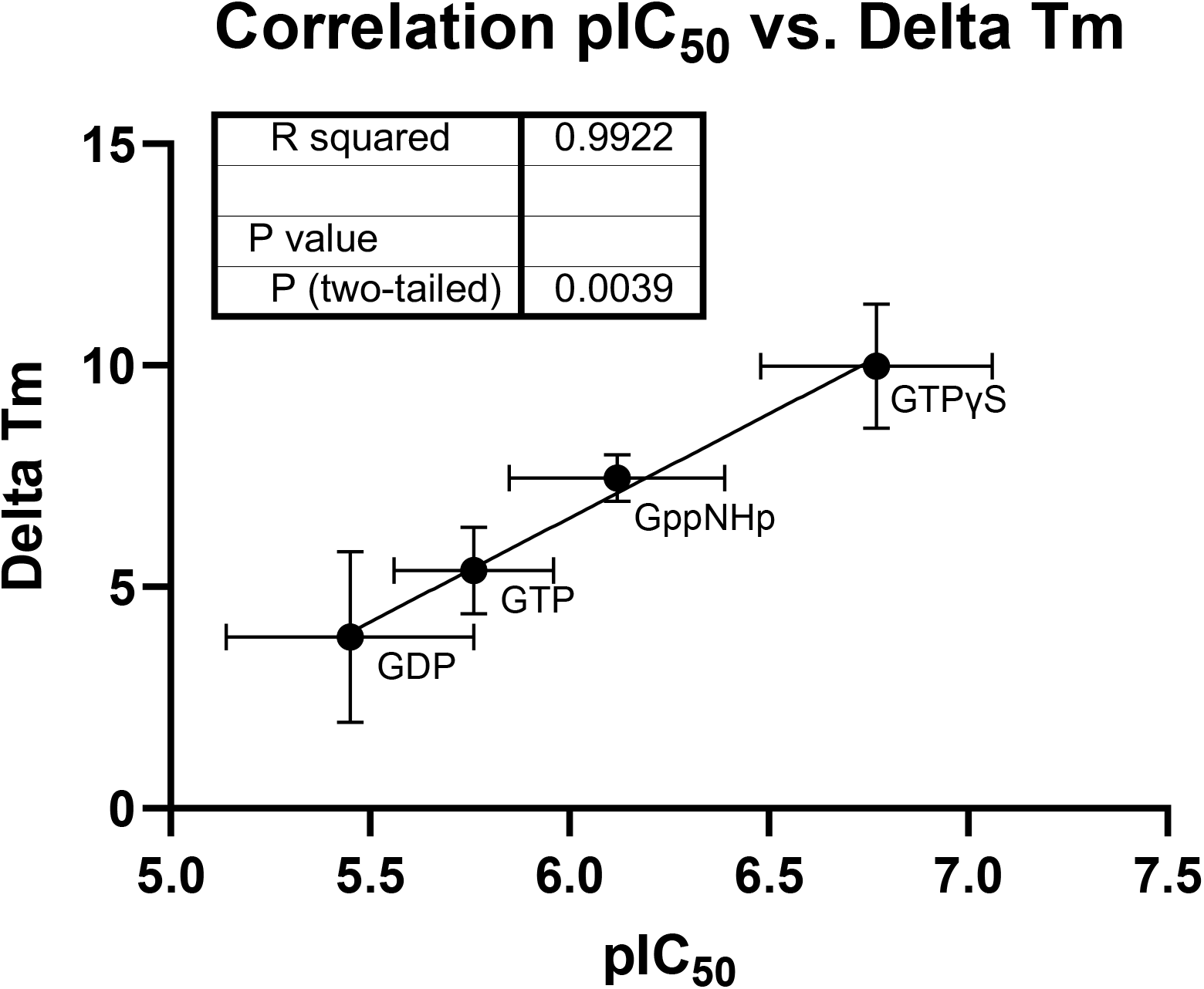
Correlation between SolThermoBRET derived T_m_’s determined in the presence of high ligand concentrations and IC_50_ determinations using Isothermal SolThermoBRET. Data shown as mean ±standard error (n≥3).

## Discussion

Drug research is an intricate and resource-intensive process, requiring significant human effort and material resources at every stage. One early stage in this research and development process, which is pivotal to compound progression, is confirmation of target engagement that gives confidence to medicinal chemists to iteratively improve the design of their drugs [22].

The current investigation into ligand induced thermostability of the Gα_i1_ protein is crucial for understanding the utility of the ThermoBRET method, as a reporter of target engagement, when applied to soluble proteins. As a first step in this process, determining a suitable concentration of the Gα_i1_ protein, labelled with tsNLuc, proved crucial and fundamental for reliable results. The luminescent levels at various dilutions of the unpurified protein were carefully examined, ensuring that the subsequent experiments were conducted under conditions reflective of the protein’s native state and using a concentration of protein at which the tsNLuc substrate furimazine, would not be significantly depleted. This optimization step is illustrated in Figure 2 and underscores the efficiency of the SolThermoBRET assay in comparison to traditional biophysical techniques which traditionally require micro-milligrams of purified protein [23, 24].

Importantly ligand-induced stabilisation of the protein was investigated using non-purified samples. By monitoring the unfolding of the protein in the presence of a fixed concentration of the different nucleotides/sides and SYPRO Red, we gained valuable insights into the impact of various ligands on the protein’s structural integrity. The results, depicted in Figure 3, highlighted the order of ligand-induced thermostability shifts, with GDP, GTP, GppNHp, and GTPγS all affecting the protein in a distinct manner. Notably, the absence of a shift with ADP suggests either a lack of binding to the protein or an affinity below the employed concentration.

Our observations regarding ligand-induced thermostability shifts align with previous studies utilizing biophysical techniques. The order of ligand effects on the protein stability (GDP < GTP < GppNHp < GTPγS), as demonstrated in Figure 3, is consistent with other reports in the literature [25]. This conformity strengthens the validity of our results and establishes a foundation for cross-study comparisons.

Isothermal SolThermoBRET experiments were conducted to quantitatively measure the relative affinity of the different nucleotides/sides tested. The inhibitory effects of ligands on protein unfolding at a constant temperature, are shown in Figure 4A-D. The IC_50_ values measured for these ligands (see Table 2) provide a quantitative measure of the concentration at which half-maximal inhibition occurred, offering a valuable metric for comparing ligand affinities. The observed order of ligand affinities (GDP < GTP < GppNHp < GTPγS) is consistent with trends reported in analogous studies studying displacement of high affinity agonist binding, providing a basis for understanding the relative binding strengths of nucleotides/sides to the soluble protein [26, 27].

The correlation analysis depicted in Figure 5, demonstrating a strong agreement between SolThermoBRET-derived ΔTm and IC_50_ values, aligns with similar correlation analyses conducted in the literature [19]. This consistency underscores the reliability of our assay and reaffirms the relationship between ligand-induced shifts in melting temperature and their inhibitory concentrations. Such concordance with established methodologies adds weight to the robustness of our experimental design and results.

The integration of BRET technology with SYPRO Red provides a powerful approach for studying the thermostability of soluble proteins, especially those labelled with tsNLuc. This methodology allows for a comprehensive understanding of ligand-induced effects on protein stability, offering both qualitative and quantitative insights. Comparisons with findings from similar methods employed in the literature further contextualize the significance of our results.

To conclude SolThermoBRET represents a distinct nanoscale method for the detection of soluble protein thermostability being both rapid and cost-effective. SolThermoBRET not only accommodates the complex milieu of soluble proteins and essential cofactors, but also attains unprecedented sensitivity. This method is ideal for hit conformation studies and the identification of novel ligands targeting soluble proteins.

## Acknowledgements

PH is funded by an international fellowship awarded by the Office of Educational Affairs Thailand.

